# Combinatorial Protein Language Model–Guided Engineering of TEV Protease for Enhanced Stability and Production

**DOI:** 10.64898/2025.12.19.695604

**Authors:** Tek-Hyung Lee, Songming Liu, Yifan Li, Trang Nguyen, Zhaoren He, Sagar Khare, Li Yi

## Abstract

Industrial enzyme engineering focuses on improvement of enzyme production yield, stability, catalytic activity, and substrate specificity, but often suffers from low efficiency with time-consuming and labor-intensive design and screening processes of massive libraries. Recent advances in AI and machine learning created protein language models trained by numerous datasets and shed new lights to speed up the enzyme engineering processes with high accuracy structural prediction. Here, we developed a highly efficient enzyme engineering strategy combining three protein language models (xTrimoMPNN-Thermo, ESM-IF, and MPNNsol) and use it to generate TEV protease variants with improved expression, stability, and function. The results indicated that a small number of TEV protease designs (<50 designs) were sufficient to develop variants with desired properties, demonstrating its high efficiency. Our strategy could be broadly applied to accelerate designing and engineering various industrial enzymes.

## Introduction

Enzymes are industrially attractive biological catalysts, especially in food and pharmaceutical sectors, since they can catalyze specific reactions rapidly and selectively to generate desired products, significantly lowering the reaction energy barrier. However, industrial application requires high production yield, stability, and activity of enzymes that can sustain under harsh process conditions, which is often challenging to achieve because in many cases, enzymes are aggregated, denatured, or lose their catalytic activity.

There have been numerous research efforts to engineer enzymes to overcome those hurdles. Directed evolution and rational design engineering have shown remarkable successes, becoming leading strategies in enzyme engineering^1,2^. However, directed evolution often suffers from time-consuming and labor-intensive processes to construct and test numerous libraries. Rational design strategy can reduce the mutational space aiming specific regions, but it requires a deep understanding of enzyme structures and functional mechanisms to select target sites for engineering, which is not always available. With recent advances in AI and machine learning, high accuracy structural prediction methods such as AlphaFold and Rosettafold and protein language models (pLM) have been developed, learning underlying patterns of proteins sequences from massive protein databases, which significantly changed enzyme engineering methodologies. These pLMs accelerated the design of novel enzymes which are largely different from evolved natural proteins, dramatically reducing vast mutational spaces for experimental testing^3,4^.

Inspired by recent progress, we employed multiple protein language models trained upon thermostable protein databases to develop a highly efficient enzyme engineering strategy to improve its production and stability. Using TEV protease as the example, three rounds of in silico design were performed with optimized filtering, generating a small set of variants (< 50) for experimental validation. Most TEV protease variants exhibited much higher stability and production yield compared to the parent TEV protease (TEVp-S219D). The TEV protease variants with high activity were further characterized, indicating that these variants showed improved turnover numbers with intact substrate specificity.

## Results

### Designing strategy of TEV protease variants

To design high stability TEV protease variants *in silico*, an inverse folding model, called xTrimoMPNN-Thermo, was derived from ByProt Non-Autoregressive ProteinMPNN architecture^5^, training it with two thermostability datasets (learn2thermDB and Meltome^6,7^) (**Figure 1A**). To evaluate thermostability of designed TEV protease variants, xTrimoTm3b and xTrimoOGT3b, two thermostability predicting models were used.

**Figure 1:**
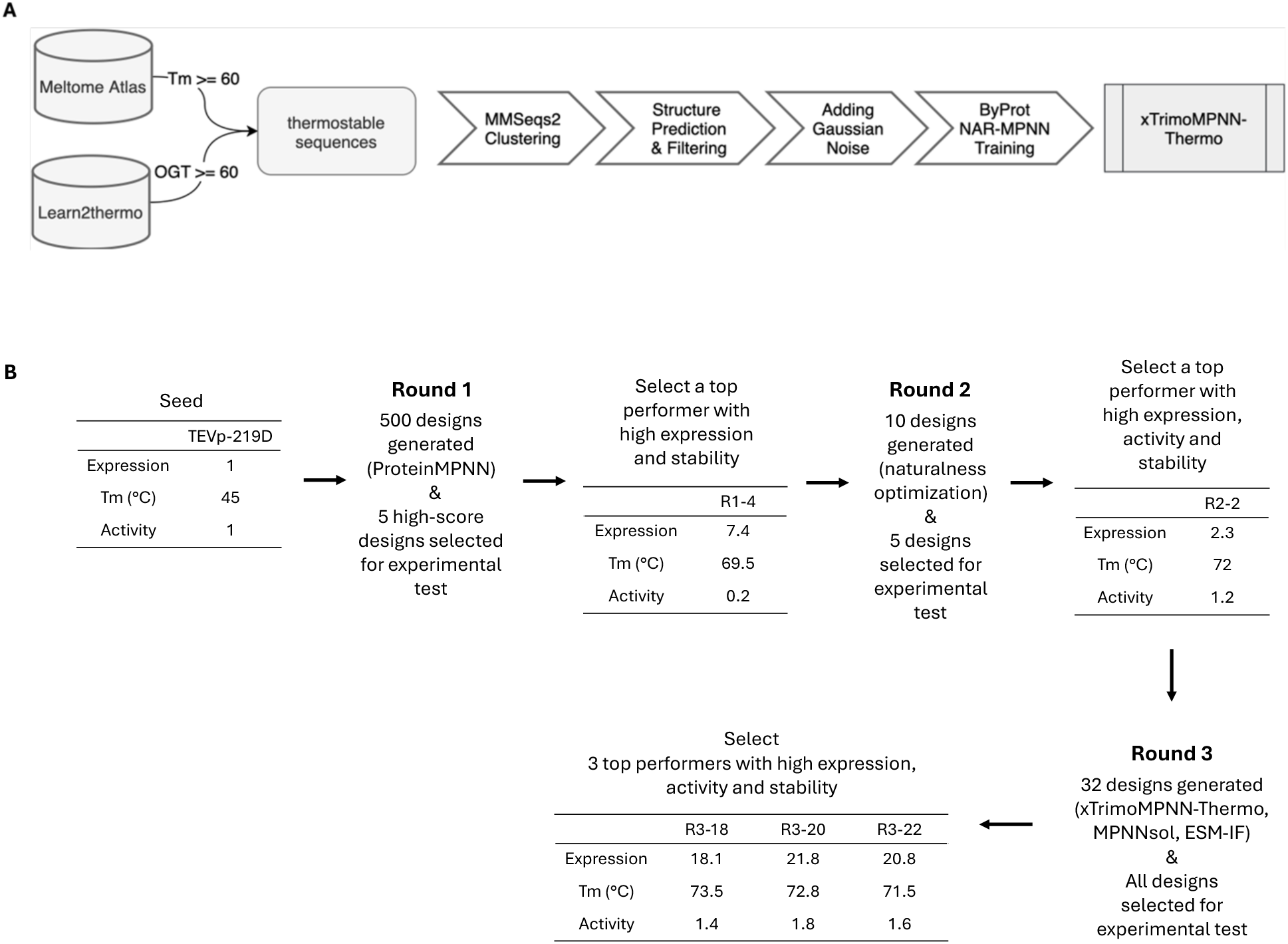
Flow chart of computational design strategies of de novo TEV protease engineering. (A) Schematic of the strategy to generate sequences of thermostable TEV protease variants. Thermostable protein sequences were collected from two databases (Meltome and Learn2thermo)^6,7^, and the predicted structures were fine-tuned using the ByProt non-autoregressive ProteinMPNN^5^. (B) Three rounds of computational design and experimental characterization to generate TEV protease variants with high stability, expression, and catalytic activity

Three-round optimization processes were performed to generate high stability TEV protease variants (**Figure 1B**). Firstly, TEVp-S219D, a TEV protease with S219D mutation (PDB ID: 1LVM) was used as the structural input of original ProteinMPNN^3^, fixing the catalytically active sites and 50% of the most conserved residues to preserve catalytic activity^4^. 500 designs were generated, and 5 variants (R1 variants) with high predicted melting temperature (xTrimoTm3b prediction) and high predicted thermostability improvement (xTrimoOGT3b prediction) were selected through in-silico filtering (**Table 1**). Production yield and activities of the 5 variants were evaluated, and the highest production variant (R1-4) was selected (**Supplementary Figure 1**) to further optimize its sequence naturalness for lowering the possibility of potential aggregation or poor stability. 5 new variants were designed in the second round, sequentially introducing additional^8^ (**Figure 2A**, **Supplementary Figure 2**). After characterization, R2-2 exhibited high activity and thermostability, which was selected for the third-round optimization (**Supplementary Figure 1**).

**Figure 2.**
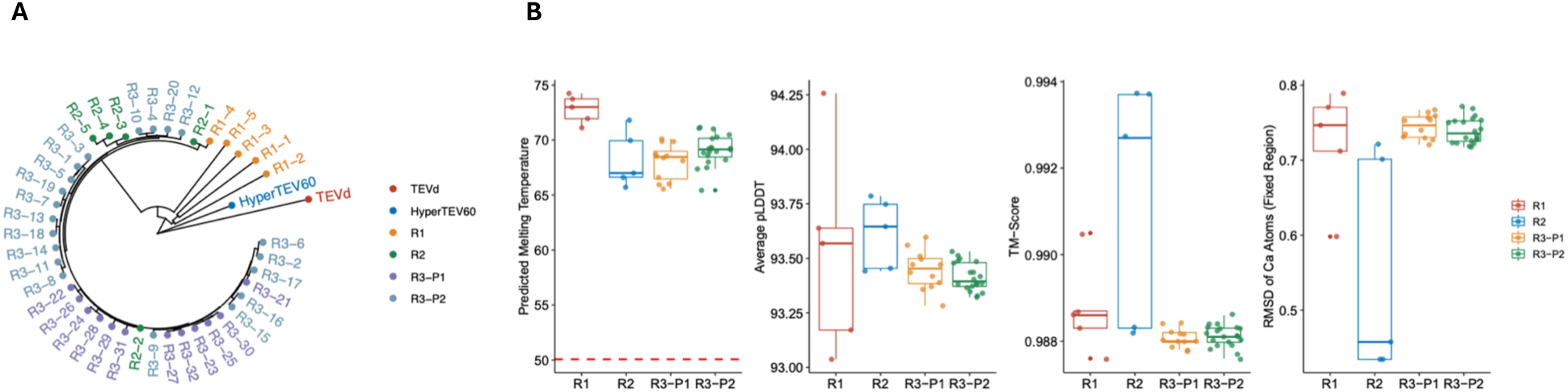
*In silico* designs of TEV protease enzyme using three pLM models with naturalness optimization. (A) The phylogenetic tree of all R1/R2/R3 designs utilizing the edit distance. (B) *In silico* model prediction score plots (whisker plot), for melting temperature predicted by xTrimoTm3b model, average pLDDT, TM scores, and Ca atom RMSD in fixed regions (from left to right). The red dashed line indicates the melting temperature of TEVp-S219D. R3-P1 denotes R3 Paradigm 1 (designed using a xTrimoMPNN-Thermo model only), and R3-P2 denotes R3 Paradigm 2 (designed using xTrimoMPNN-Thermo,MPNNsol, and ESM-IF).

**Table 1.** pLM prediction scores for R1, R2, and R3 TEV protease designs.

| Variant Group | Predicted Melting Temperature (xTrimoTm3b) | Stability Improvement (xTrimoOGT 3b) | Average pLDDT | TM Score | RMSD<sub>Ca</sub> | RMSD<sub>Ca</sub> (Fixed Region) |
| --- | --- | --- | --- | --- | --- | --- |
| R1 | 72.8 ± 1.3 | 12.1 ± 0.5 | 93.5 ± 0.5 | 0.99 ± 0.001 | 0.8 ± 0.1 | 0.7 ± 0.1 |
| R-2 | 68.2 ± 2.6 | 14.0 ± 1.2 | 93.6 ± 0.2 | 0.99 ± 0.003 | 0.6 ± 0.2 | 0.5 ± 0.2 |
| R3-P1* | 68.0 ± 1.6 | 12.8 ± 0.8 | 93.4 ± 0.1 | 0.99 ± 0.001 | 0.9 ± 0.01 | 0.7 ± 0.02 |
| R3-P2* | 69.1 ± 1.4 | 13.8 ± 1.0 | 93.4 ± 0.1 | 0.99 ± 0.001 | 0.9 ± 0.01 | 0.7 ± 0.02 |

R3-P1\* and R3-P2\* denote R3 paradigm 1 designs and R3 paradigm 2 designs, respectively.

In the third-round optimization, two selection paradigms were applied to prioritize the substitutions benefiting enzyme stability. The first one (paradigm 1) relied on a single-model criterion, ranking substitutions solely by newly trained xTrimoMPNN-Thermo. The second one (paradigm 2) enforced a consensus requirement: a candidate substitution was retained only if it has a higher score exceeding that of the wild-type residue simultaneously across three models: xTrimoMPNN-Thermo, MPNNsol^9^, and ESM-IF. Relative to paradigm 1, paradigm 2 yielded a more conservative mutation set and likely reduced model-specific idiosyncrasies. Finally, a total of 32 variants, including 12 from paradigm 1 and 20 from paradigm 2, were selected in the third round for experimental validation (**Figure 2A**, **Supplementary Table 1**).

Vigorous filtering criteria (predicted Tm ≥ 60, average pLDDT ≥ 90, TM-score ≥ 0.9, and C_α_ RMSD ≤ 1) were applied for three-round optimization. The predicted model scores for R1, R2, and R3 designs were confirmed to comply with the criteria before experimental validation (**Figure 2B**).

### TEV protease designs exhibit high production yields, thermostability, and catalytic activities

The expression of the R3 designs was evaluated in *E.coli*. All 32 designs were expressed in a soluble form, and their expression levels were significantly higher (5-25 times) than TEVp-S219D (**Figure 3A, Supplementary Figure 3 and 4A**). The top 6 high-production variants showed similar expression levels compared to a recently published TEV variant, HyperTEV60^4^. These results confirmed that the pLM-based designs effectively enhanced heterologous enzyme production level, possibly increasing its folding efficiency.

**Figure 3.**
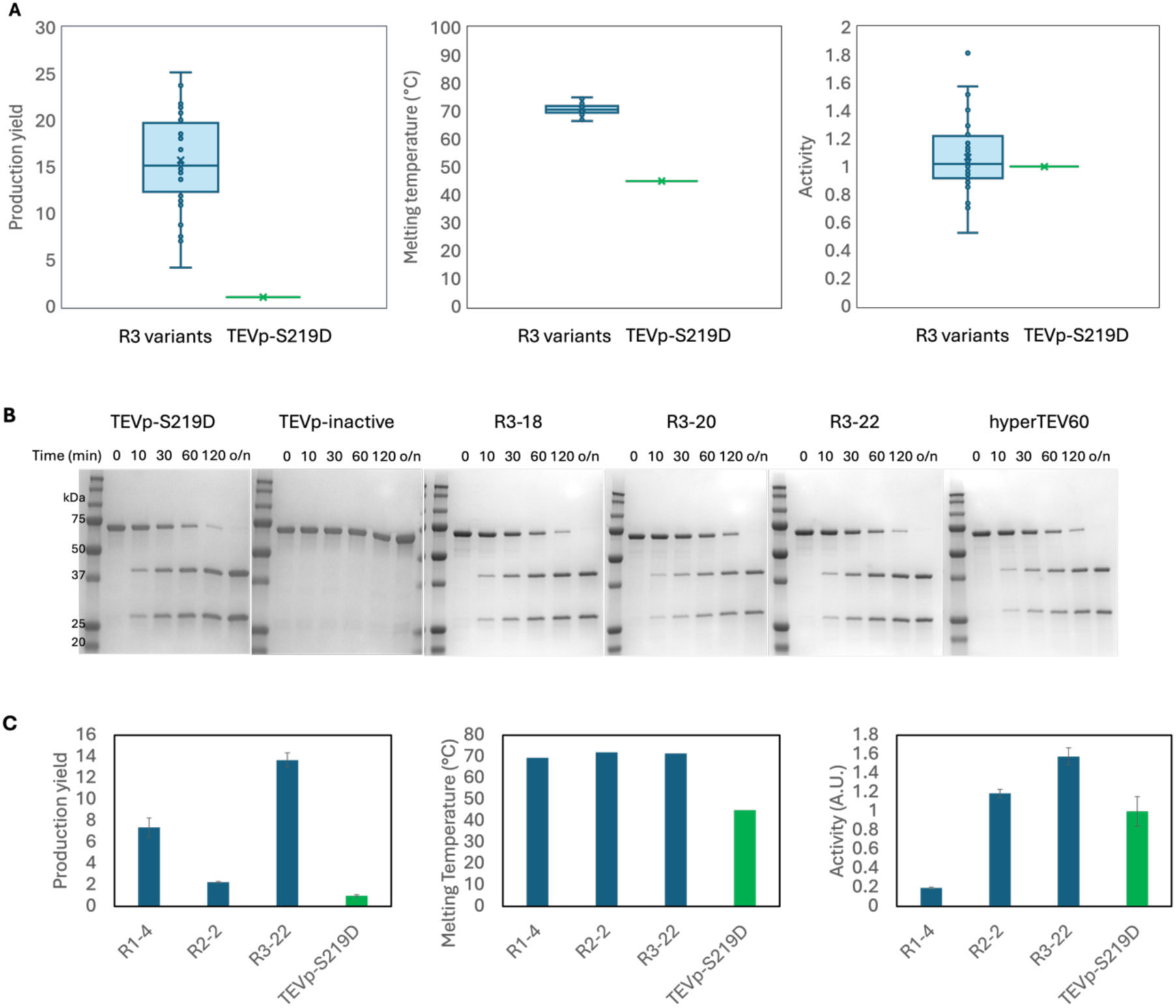
The characterization of production yield, stability, and activity of TEV protease designs. (A) The characterization of production yield of TEV protease designs in *E.coli* compared to control TEV proteases. The production yields are normalized by the production yield of TEVp-S219D. The evaluation of thermostability of R3 TEV protease designs compared to control TEV proteases. The characterization of catalytic activities of R3 TEV protease designs compared to control TEV proteases using peptide substrates. Reactions were carried out by mixing 1mg (∼ 0.4 mM) TEV protease with 0.25 mM peptide substrates, which were incubated at 37°C for 1 hour. The results were quantitated by fluorescence intensity change and normalized by TEVp-S219D. All data points were plotted using a whisker plot where the mean value was marked as “X.” (B) The characterization of catalytic activities of R3 TEV protease designs compared to control TEV proteases using protein substrates (MBP-ENLYFQ↓S-EGFP). Reactions were carried out by mixing 0.5mg protease with 50mg protein substrates, which were incubated at 30°C for 0 min, 10 min, 30 min, 1 hour, 2 hours, and overnight (18 hours). The proteolytic progression was monitored by SDS-PAGE. (C) The plots of production yield, thermostability, and catalytic activity of high-performance designs of R1, R2 and R3. The production yields are normalized by the production yield of TEVp-S219D. The catalytic activities are also normalized by TEVp-S219D activity.

Next, the thermal shift assay was performed to evaluate the thermostability of the R3 designs. All variants showed elevated melting temperatures (Tm, 69°C - 75°C) higher than that of TEVp-S219D (45 °C). Their Tm values were comparable to the Tm (75 °C) of HyperTEV60, verifying successful stability improvement (**Figure 3A, Supplementary Figure 4B**). It is also worth pointing out that the experimentally measured Tm values showed a good correlation with predicted Tm values (**Supplementary Figure 5**).

Subsequently, the catalytic activities of R3 designs were quantitated using a 5-FAM/QXL 520 peptide substrate (AnaSpec). All 32 R3 designs generated high fluorescence signals upon substrate cleavage after 60 min at 37°C, showing activities comparable to or higher than TEVp-S219D (**Figure 3A, Supplementary Figure 4C**). Interestingly, 8 out of 32 designs presented more than 20% increased activity than TEVp-S219D. Combining the production, Tm, and activity pre-screening results, three designs (R3-18, R3-20, and R3-22) were further examined of their catalytic activities against an MBP-EGFP fusion protein substrate, where the TEV protease recognition site (ENLYFQ↓S) was inserted between MBP and EGFP. Substrate cleavage progression was evaluated by SDS-PAGE and all three variants showed that they preserve high cleavage activity comparable to that of TEVp-S219D. Including HyperTEV60, all variants showed complete digestion after 18 hours incubation at 30°C with 1:100 ratio of enzyme : protein substrate. (**Figure 3B**).

R3-18, R3-20, and R3-22 designs presented good biochemical properties. A detailed kinetic characterization using 5-FAM/QXL 520 peptide substrate showed that R3-20 and R3-22 are faster protease with k_cat_/K_m_ of 13.3±0.1×10^−4^ µM^−1^ s^−1^ and 14.4±0.3×10^−4^ µM^−1^ s^−1^, respectively. Comparably, k_cat_/K_m_ of TEVp-S219D is 9.3±0.8×10^−4^ µM^−1^ s^−1^ and R3-18 design is 5.9±0.1×10^−4^ µM^−1^ s^−1^ (**Table 2, Supplementary Figure 6**). All TEV protease variants have comparable K_m_ values, while R3-20 and R3-22 designs showed higher turnover numbers (k_cat_) of 32.9 ± 2.7 ×10^−4^ s^−1^ and 31.6 ± 4.7 ×10^−4^ s^−1^, respectively.

**Table 2.** Kinetic parameters of top three R3 TEV protease variants.

|  | TEVp-S219D | R3-18 | R3-20 | R3-22 | HyperTEV60 |
| --- | --- | --- | --- | --- | --- |
| K<sub>m</sub> (μM) | 2.0 ± 0.2 | 2.0 ± 0.2 | 2.5 ± 0.2 | 2.2 ± 0.4 | 2.1 ± 0.1 |
| k<sub>cat</sub> (x10<sup>-4</sup> s<sup>-1</sup>) | 18.7 ± 0.1 | 11.8 ± 1.4 | 32.9 ± 2.7 | 31.6 ± 4.7 | 7.2 ± 0.3 |
| k<sub>cat</sub>/K<sub>m</sub> (x10<sup>-4</sup> μM<sup>-1</sup> s<sup>-1</sup>) | 9.3 ± 0.8 | 5.9 ± 0.1 | 13.4 ± 0.1 | 14.4 ± 0.3 | 3.5 ± 0.1 |

In addition, high-performance designs from each round (R1-4, R2-6, and R3-22) were compared with TEVp-S219D. All variants showed comparable thermostability, which is significantly improved compared to the TEVp-S219D (**Figure 3C**). The R3-22 exhibited the highest production yield and catalytic activities. R1-4 showed significantly higher production yield compared to TEVp-S219D, but its activity was lower. Interestingly, the production yield of R2-6 was lower than R1-4, while its activity was higher, suggesting a possible trade-off between expression and activity during naturalness optimization. Taken together, three rounds of design optimization with three pLMs successfully enhanced TEV protease stability, activity and production yield.

## Discussion

In this study, we pursued the high efficiency design process of enzyme engineering by employing three protein language models. The design and optimization strategy presented here is highly efficient to develop active enzymes with improved production yield and stability, given that only a small number of variants (42 variants) were tested. Among them, 37 TEV protease variants (one R1, four R2 designs and 32 R3 designs) exhibited significantly improved production yield and stability compared to the parent TEVp-S219D, preserving its original activity or exhibiting enhanced activities. We reasoned this improvement from 1) design using a protein language model trained with large scale dataset of thermostable proteins, 2) stringent filtering process using xTrimoOGT3b and xTrimoTm3b, 3) multiple rounds of optimization, and 4) fixing highly conserved residues including active site residues to maintain enzyme activities.

After detailed characterization, two designs, R3-20 and R3-22, were identified to be ∼ 50% faster than the parent TEVp-S219D, with > 20-fold of production yield, and improved Tm from 45 °C to 72 °C. The sequence similarity of R3-20 and R3-22 against TEVp-S219D is about 70% (**Supplementary Table 1**). It needs to be pointed out that a recent study on TEVp-S219D showed that fixing less numbers of residues (<30%) significantly lowered the success rate of generating active enzymes^4^. Therefore, further studies remain to evaluate if design and optimization strategies can improve success rate for designing a low sequency identity enzyme.

A similar strategy was utilized in R1 designs of our studies compared to the recently published study^4^, in which MPNN was applied to significantly increase the production yield, stability, and activity of the TEVp-S219D. While the recent study generated designs with fixing the active site and various numbers of most conserved residues (0, 30%, 50%, and 70%) in TEV protease family, here we designed TEV protease variants, fixing the active site and the top 50% most conserved residues, and using the latest version of ProteinMPNN (git commit version: 8907e66) with higher temperature (T=0.7) to obtain more diverse designs. We further enhanced naturalness (R2 designs) and applied three models (xTrimoMPNN-Thermo, MPNNsol, and ESM-IF), which improved production yield and activity (R3 designs) (**Figure 3C)**.

In our studies, R3-20 and R3-22 designs showed moderately better activity than HyperTEV60. We speculate that this discrepancy is caused by different substrates being used to screen TEV protease variants in two studies. Although P1’ substrate specificity was always neglected in TEV protease, its lower proteolytic cleavage against bulky residue at P1’ site, such as Trp, was previously reported^10,11^. R3-20 and R3-22 designs exhibited high catalytic activities comparable to both TEVp-S219D and hyperTEV60 for substrates containing small residues at P1’ site, such as ENLYFQ↓G and ENLYFQ↓S (**Figure 3B**). However, when the P1’ site was replaced by bulky hydrophobic coumarin ring, TEVp-S219D as well as R3-20 and R3-22 designs showed significant loss of proteolytic activities compared to HyperTEV60 (**Supplementary Figure 6**). It is hard to interpret this discrepancy intuitively, because sequences around the active site were identical between R3-20 and R3-22 designs, hyperTEV60 and TEVp-S219D, and their predicted structures are well-aligned. Considering approximately 20% of residues are changed in hyperTEV60, R3-20 and R3-22 designs, some distant residues might cause their different substrate tolerance at the P1’ position, as a recent study reported that the mutation of R203, which is located far from the catalytic center contributes to providing positive electrostatic surface patch, thus affecting the protease’s P1’ substrate specificity^10^.

## Supporting information

Supplementary Sheet

## Supplementary Information

### Supplementary Methods

#### 1. Computational methods

##### 1.1 Thermostable model training

Data were obtained from two thermostability dataset: learn2thermDB, a large dataset consisting of 24M protein sequences with optimal growth temperature (OGT) and 69M protein pairs based on homology^6^, and Meltome, a thermal stability atlas of 48,000 proteins across 13 species ranging from archaea to humans and covering melting temperatures (Tm) of 30–90 °C^7^.

For the xTrimoMPNN-Thermo model training, the sequences from learn2thermDB with OGT ≥ 60°C and sequences from Meltome atlas with melting temperature ≥ 60°C were preserved (N=387,255). Those sequences were clustered using MMseqs2 easy-cluster module with an identity set to 0.4 and sequence length being limited to 50∼500 aa^12^. A total of 99,275 thermostable sequences were obtained, and their structures were predicted using xt-Fold model^8^. A total of 16,675 sequences with high-confidence structures (min plddt >= 60, median plddt >= 90, and mean plddt >= 90) were selected and a gaussian noise (std=0.05Å) were added to their backbones^3^. Finally, the xTrimoMPNN-Thermo model was trained on ByProt non-autoregressive (NAR) ProteinMPNN architecture utilizing those noised backbones^5^.

To evaluate the model thermostability, two regressive models based on xTrimoProtein model were fine-tuned. One is xTrimoOGT3b, which was trained using learn2themoDB homologous sequence pairs (identity >= 0.8) and using the difference of OGT as prediction targets; the other is xTrimoTm3b, which was trained using Meltome atlas data and the Tm as prediction targets.

##### 1.2 Thermostable TEV design protocol

The crystal structure of TEVp-S219D (PDB ID: 1LVM) was obtained from the Protein Data Bank. A three-round procedure was then performed to generate designs for experimental characterization.

###### Round 1

Firstly, chain A (position 1-221) of TEVp-S219D structure was extracted and used as the structural input for ProteinMPNN. Active sites and the 50% most conserved residues were fixed following the protocol established by Sumida et al^4^. A total of 500 designs were generated, and the top 5 designs (R1 designs) after in-silico filtering were selected for validation.

###### Round 2

After the experimental validation, the R1-4 design, which showed the highest production yield and activity, was selected for further optimization based on the naturalness score. Specifically, the naturalness score was defined as the average token probability of the current sequence based on crystal structure using ESM-IF model^9^. ESM-IF model was iteratively run for 10 times to obtain the probability matrix (sequence length (L) x 20 canonical amino acids), introducing one mutation each time by replacing the position in current sequence with lowest token probability to the amino acid with highest probability at corresponding position (**Supplementary Table 1**). 5 designs (R2 designs) from iteration step 2, 4, 6, 8, and 10, were selected for experimental validation.

###### Round 3

The sequence R2-2 was selected as the seed to further improve enzymatic stability. Two complementary design strategies were implemented. In the first strategy, the xTrimoMPNN-Thermo model was employed to generate a set of R3 designs. Saturation mutagenesis was performed across positions (excluding the first 10 amino acids and fixed regions) of the seed sequence, and top 20 substitutions with the highest token probabilities, as predicted by the xTrimoMPNN-Thermo model were chosen. These substitutions were then manually curated, and 12 designs were generated for subsequent experimental evaluation.

In the second strategy, three models—xTrimoMPNN-Thermo, MPNNsol, and ESM-IF—were applied in combination to generate another set of R3 designs. For each residue position, only substitutions with token probabilities exceeding that of the seed (R2-2) residue in all three models were retained. From these, the first 10 residues and those in fixed regions were excluded, and high-ranking substitutions according to the xTrimoMPNN-Thermo model were selected. In addition, a subset of substitutions from the hyperTEV60 dataset^4^ that altered the same residue positions but to different amino acids was incorporated, in order to increase the probability of identifying improved variants. We finally selected 19 substitutions with manual curation.

In both strategies, substitutions reverting to the wild-type sequence were excluded. The remaining single-point substitutions were combined to construct double and triple mutants. These sequences were embedded using the xTrimoProtein model, and the resulting embeddings were subjected to t-distributed stochastic neighbor embedding (t-SNE) for dimensionality reduction and clustering into 32 groups. Representative designs from each cluster were manually selected to maximize sequence diversity while maintaining a limited total number of variants for experimental testing. Total 32 designs were selected for experimental validation.

##### 1.3 Design filtering

Structures were predicted using an in-house developed xTrimoMonomer model (unpublished) for all designs generated in Round 1 and Round 2. xTrimoTm3b model was employed to predict the melting temperature for designed sequences, and xTrimoOGT3b was used to predict thermostability improvements along with TEVp-S219D sequence as an input.

For R1 designs, 38 of 500 designs passed the in-silico filtering using the criteria: 1) xTrimoTm3b predicted melting temperature ≥ 60, and xTrimoOGT3b predicted thermostability improvements >= 10; 2) average pLDDT of the whole protein and the fixed regions ≥ 90; 3) TM-score ≥ 0.9 comparing to 1LVM structure with C_α_ RMSD of whole sequence ≤ 1 and fixed regions ≤ 1. After filtering, the top 5 designs with the highest xTrimoTm3b predictions were selected. Using a similar strategy, all 5 R2 designs were passed the filtering.

##### 1.4 Sequence Identity Calculation

All designed sequences were aligned to the TEVp-S219D sequence based on residue positions for position-wise comparison. For each pair of sequences with equal length, sequence identity was defined as the percentage of positions with identical amino acids:

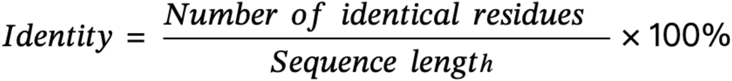

#### 2. Experimental methods

##### 2.1 Plasmid construction for TEV protease variants and their expression in *E.coli*

In-Fusion reaction (Takara Bio) was used to subclone TEV protease variant genes to pET32a (+) vector. Gene fragments (eBlock) encoding TEV protease designs were purchased from Integrated DNA Technologies (IDT). Each gene fragment contains 15 bps of flanking sequences homologous to the upstream of NdeI and downstream of HindIII of pET32a to facilitate the In-Fusion sub-cloning.

##### 2.2 TEV protease variant purification

Transformed *E.coli* cells were grown in 4 mL of autoinduction media at 37℃ until they reached an exponential phase and TEV protease variant’s expression was induced overnight at 20℃. Cells were lysed by incubating in a lysis buffer (B-PER, lysozyme, DNaseI, 2.5 mM MgCl_2_, 0.2 mM CaCl_2_, protease inhibitor cocktail, 1mM PMSF, and 1mM DTT) for 20 min at RT. The lysate was centrifuged for 30 min at 4℃. The clear supernatant (soluble lysate) was taken and mixed with the same volume of equilibration buffer (20 mM Tris-HCl, pH 8.0 and 300 mM NaCl). Then the mixture was incubated with Ni-NTA magnetic beads pre-equilibrated with the equilibration buffer for 30 min. The magnetic beads were separated on a magnet and washed three times with wash buffer (20 mM Tris, pH 8.0, 300 mM NaCl, 25 mM imidazole). The enzymes were eluted with an elution buffer (20 mM Tris-HCl, pH 8.0, 300 mM NaCl, 540 mM Imidazole). The eluent was buffer-exchanged with 50 mM Tris-HCl, pH 8.0, 300 mM NaCl using a 3kDa MWCO of Amicon ultra-centrifugal filter (Millipore Sigma). The purity of purified TEV protease variants was checked by SDS-PAGE and their concentration was determined using Nanodrop with BSA standards.

##### 2.3 TEV protease activity assay

The catalytic activity of TEV protease variants was determined by fluorometric assay using a SensoLyte 520 TEV Protease Assay kit (AnaSpec). 1 µg of purified TEV protease variants was mixed with 0.25 µM of 5-FAM/QXL^TM^ 520 TEV substrates in 100 µL of assay buffer (AnaSpec) or reaction buffer (50 mM Tris-HCl, pH 8.0, 300 mM NaCl). The reaction mixture was incubated for 1 hour at 37°C, and the fluorescent intensity of the product (5-FAM) was measured using a plate reader at Ex 490 nm and Em 520 nm with 5-FAM standards. The actual product yield was calculated using a standard curve of fluorescent 5-FAM standards. All reactions were performed in triplicate.

For the kinetic experiment, 1.8 µg of TEV protease variants were incubated with 0.125, 0.25, 0.5, 1, 2 µM of 5-FAM/QXL^TM^ 520 TEV substrates in 100 µL of reaction buffer, and fluorescent intensity was monitored every 1 min for 2 hours. Initial reaction rate was calculated for the first 10 min and used in Michaelis Menten kinetics model fitting to determine kinetic parameters (K_M_, k_cat_, and k_cat_/K_M_) by linear regression. All reactions were performed in triplicate.

##### 2.4 Protein substrate cleavage by TEV protease variants

A similar EGFP-ENLYFQ↓S-MBP fusion protein was constructed as the protein substrate^4^, but here without FKBP. 0.5 µg of TEV protease variants were mixed with 50 µg of substrate (1:100 ratio) in the reaction buffer (50 mM Tris-HCl, pH 8.0, 300 mM NaCl, 1 mM DTT) and incubated at 30°C. Time-course substrate protein cleavage was characterized by SDS-PAGE, with checking points of 0 min, 10 min, 30 min, 1 hour, 2 hours, and overnight (18 hours). The SDS-PAGE gel was stained with InstantBlue Coomassie Staining solution (Abcam) followed by being imaged using iBright 1500 imaging system (Invitrogen). The band intensities of the cleaved substrates were analyzed by densitometry.

##### 2.5 Thermal shift assay of TEV protease variants

The stability of TEV protease variants was determined by thermal shift assay. Briefly, 10-30 µg of purified enzymes were mixed with 5x SYPRO Orange dye in 25 µL of assay buffer (50 mM Tris-HCl, pH 8.0, 300 mM NaCl). The mixture was heated from 10°C to 95°C by 0.5°C (10 sec) using CFS Real-Time PCR systems (Bio-Rad) and the fluorescence intensity of SYPRO Orange dye was measured at each temperature. The fluorescence melting curve was used to determine the melting temperature (Tm) in which the second derivative of the curve slope was zero (inflection point).

### Supplementary Figures

**Supplementary Figure 1.**
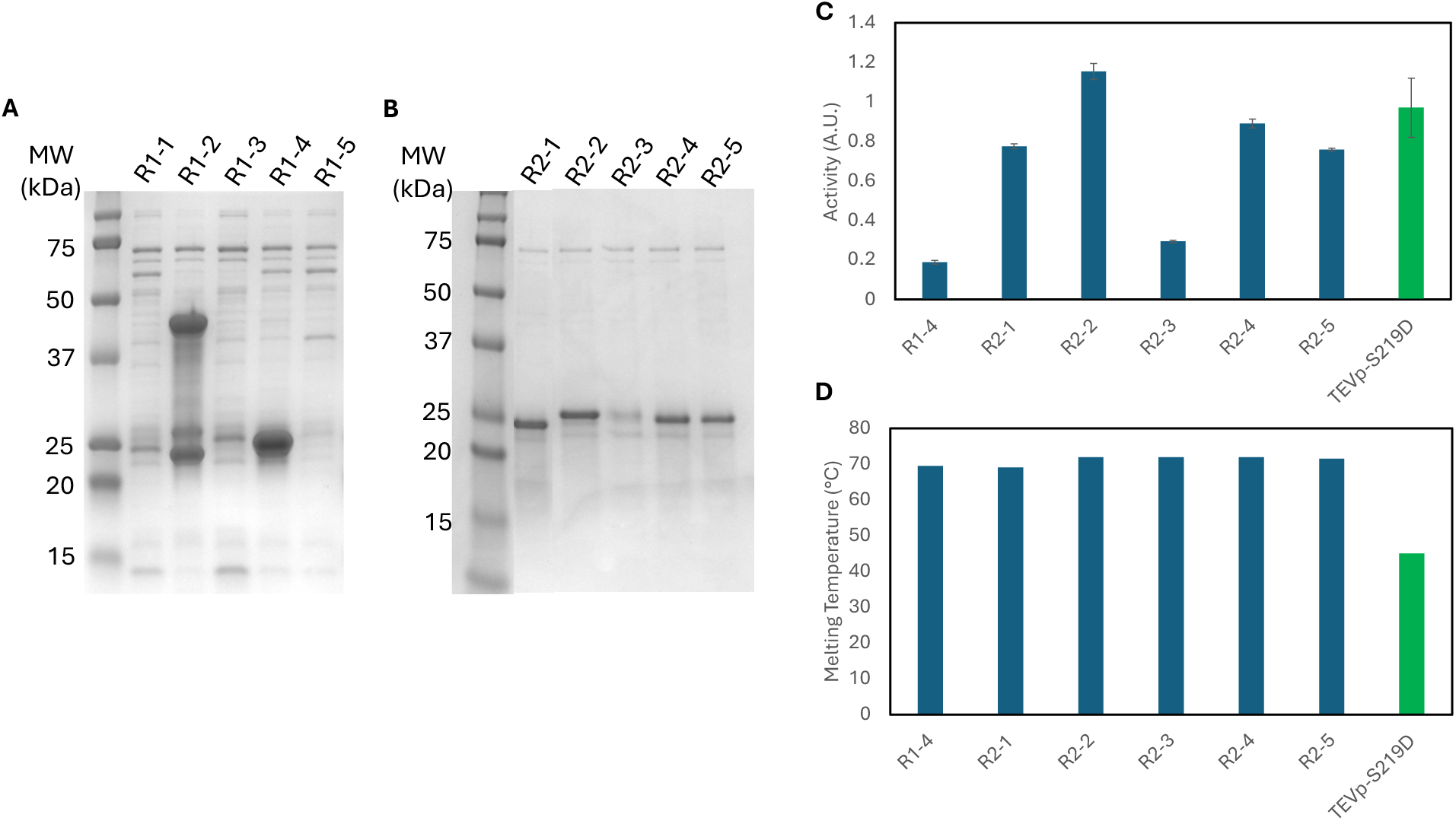
Expression, stability, and activities of R1 and R2 TEV protease variants. (A, B) 5 R1 designs and 5 R2 designs were expressed in *E.coli* BL21(DE3) and purified by Ni-NTA magnetic beads. Their expression levels and purity were verified by SDS-PAGE. R1-4 was selected for R2 designs due to its high production yield. Catalytic activities of 5 R2 designs were determined using 5-FAM/QXL^TM^ 520 TEV substrates (AnaSpec) for 1 hour at 37°C (Supplemental Methods) and compared with R1-4 and TEVp-S219D. (D) Melting temperature (Tm) of R2 designs were determined by thermal shift assay in which the fluorescence intensity of SYPRO Orange dye was measured at increasing temperatures (Supplemental Methods). The measured Tm values were compared with Tm values of R1-4 and TEVp-S219D.

**Supplementary Figure 2.**
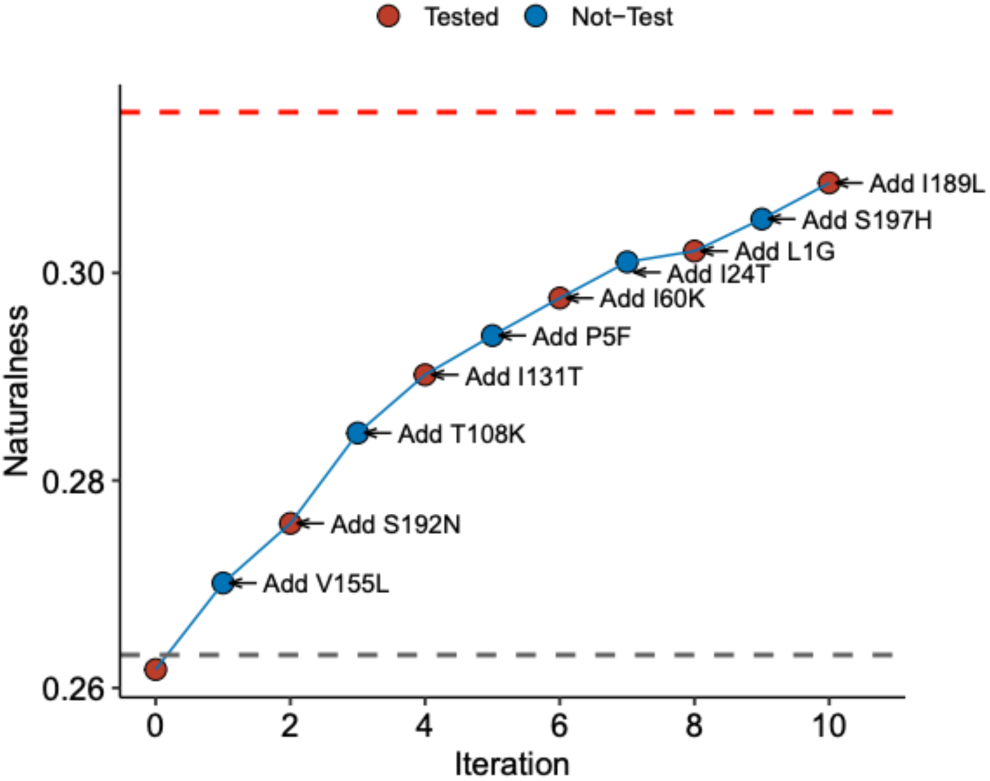
Naturalness optimization along iteration steps for R2 designs. A naturalness score plot of R2 designs, predicted by ESM-IF model. Naturalness scores were increased with iteration in which one mutation was added by replacing an amino acid with lowest token probability with an amino acid with highest probability at corresponding position (Supplemental Methods). Red and grey dashed lines indicate naturalness scores of HyperTEV60 and TEVp-S219D, respectively.

**Supplementary Figure 3.**
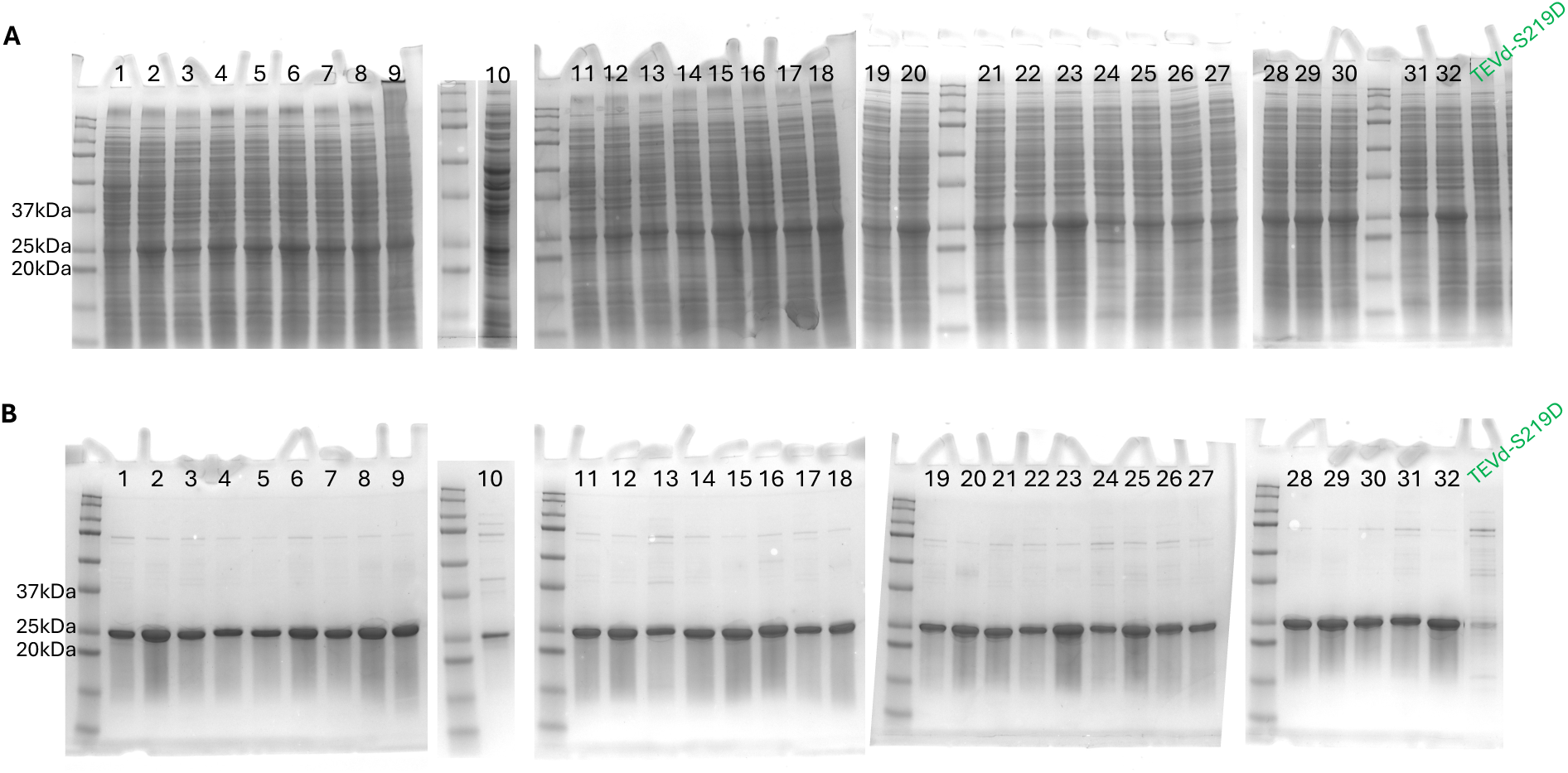
Expression and purification of R3 TEV protease variants. 32 R3 designs were expressed in *E.coli* BL21(DE3), growing them in autoinduction media. (A) The soluble fraction of TEV protease variants was extracted from *E.coli* crude lysate and their expression levels were verified by SDS-PAGE. (B) The soluble lysates were further purified by Ni-NTA affinity chromatography followed by buffer exchange and purified enzymes were confirmed by SDS-PAGE, compared with TEVp-S219D.

**Supplementary Figure 4.**
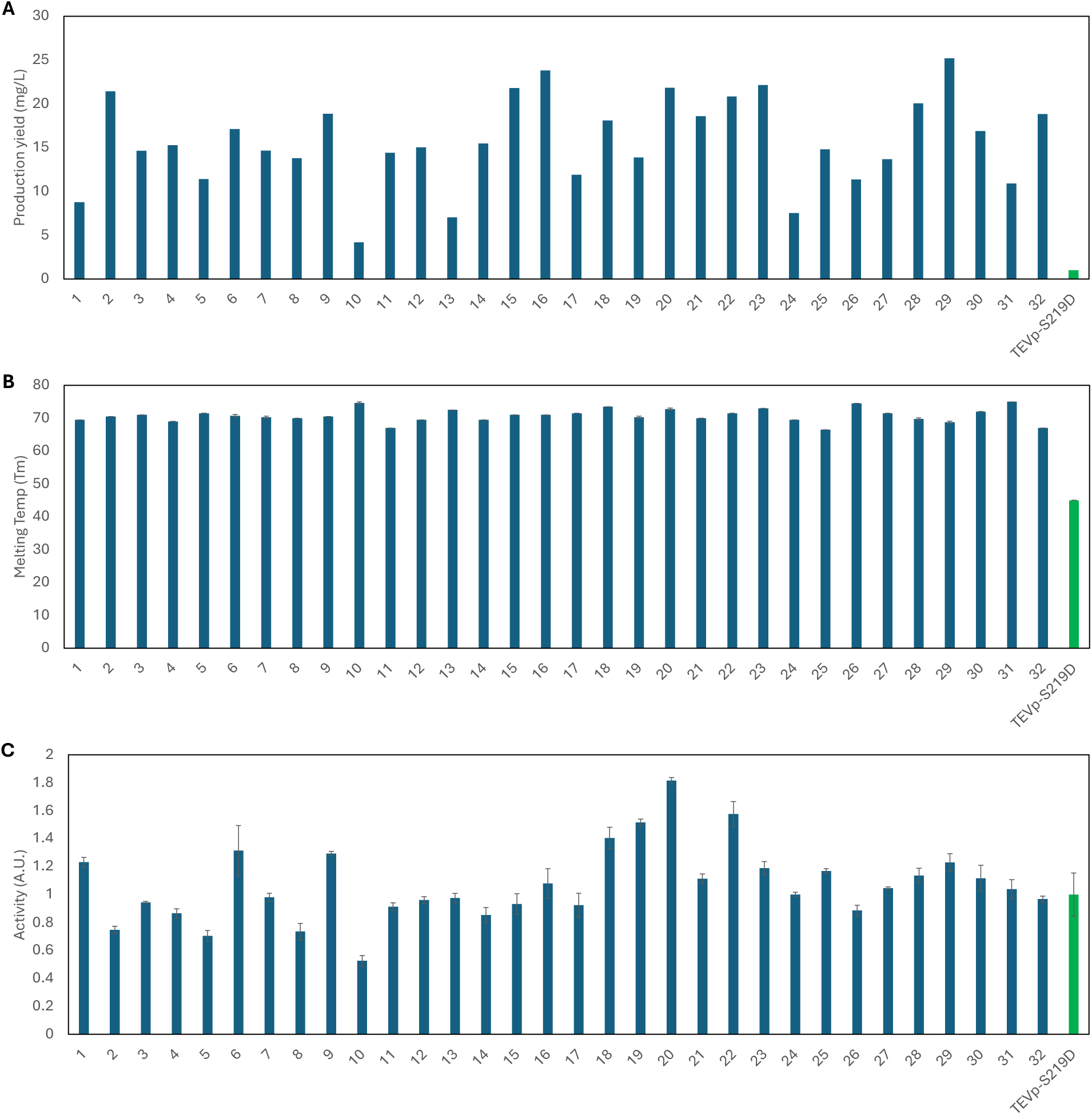
Production yield, activity and thermostability of R3 TEV protease variants. Production yield (A), thermostability (B), and catalytic activity (C), of purified 32 R3 designs were characterized (Supplemental Methods). The measured activity was normalized by the activity of TEVp-S219D.

**Supplementary Figure 5.**
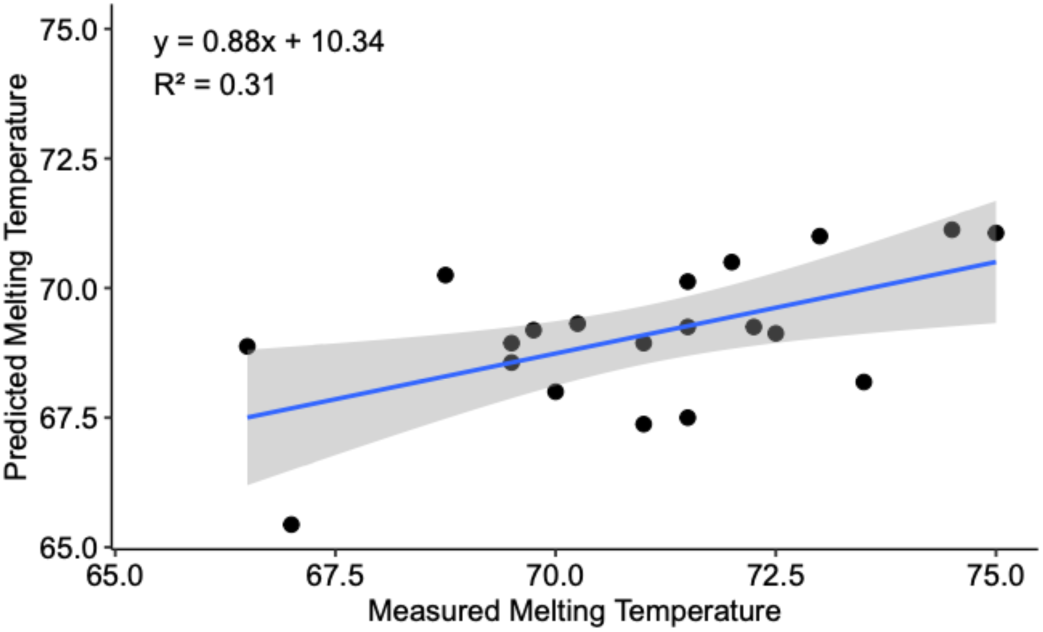
Correlation between model-predicted and experimentally measured melting temperature (Tm). The Tm values of R3 designs (paradigm 2) predicted by pLMs (xTrimoTm3b) were plotted with Tm values experimentally measured by thermal shift assay. Linear regression was used to identify a relationship between two values.

**Supplementary Figure 6.**
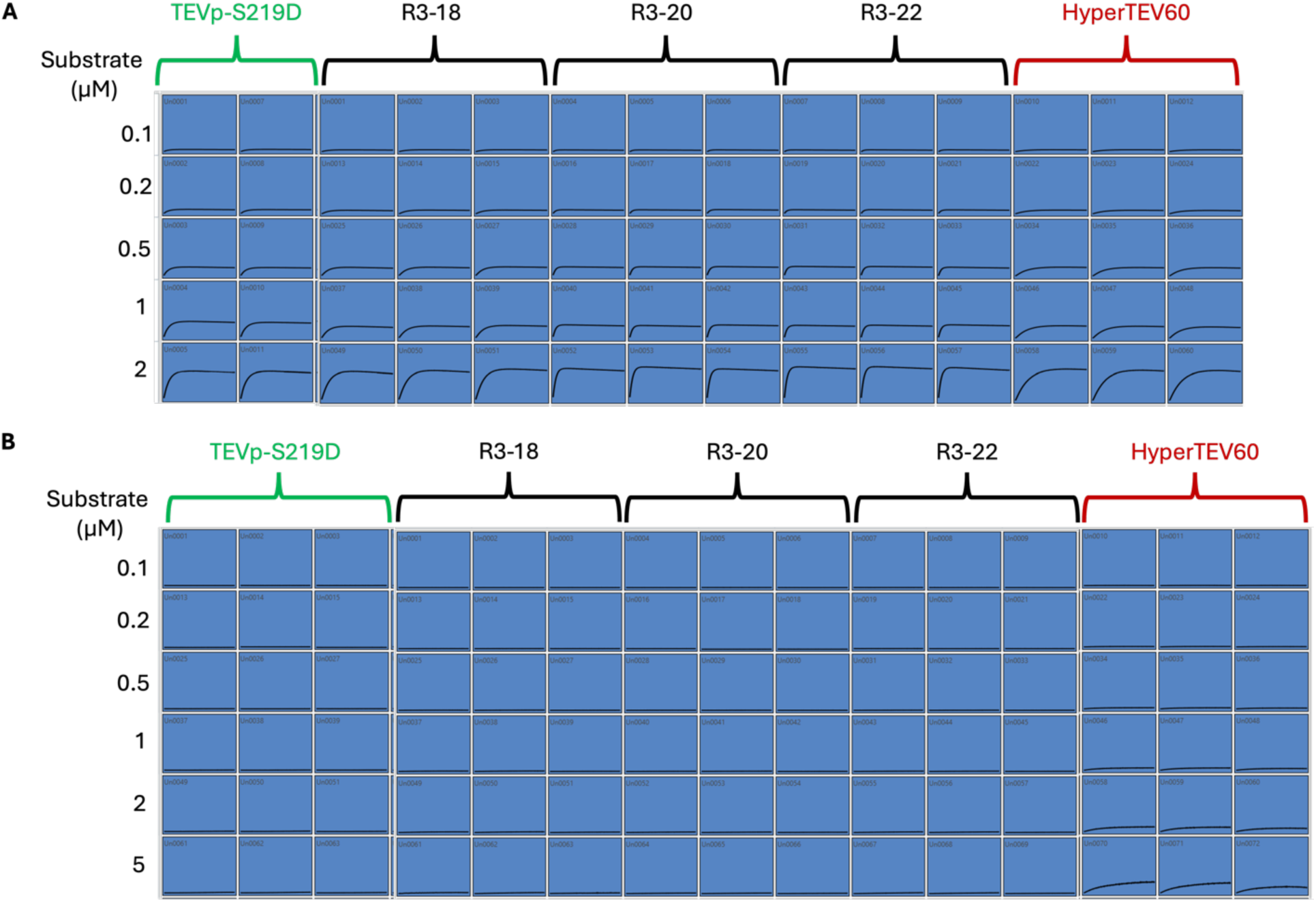
Catalytic reaction progression of TEV protease variants. 500 µM of TEV protease variants were incubated with various concentrations of substrate, and the progression of reaction was monitored by fluorescence measurement for 12 hours at 30°C. (A) Kinetics measurement for ENLYFQ↓G-containing peptide substrate (P1’ site = G) which was labeled with fluorescence dye and quencher (Anaspec) (B) Kinetics study for ENLYFQ-coumarin containing peptide substrate (P1’ site = coumarin)

### Supplementary Tables

**Supplementary Table 1.** TEV protease variants used in this study.

| TEV protease variant | Parental enzyme | Mutated residues compared to the parental enzyme | % sequence identity to TEVp-S219D | % sequence identity to hyperTEV60 |
| --- | --- | --- | --- | --- |
| TEVp-S219D<br>(PDB ID: 1LVM) | WT TEV protease | S219D (1-221 residues) | 100 | 72 |
| hyperTEV60 | TEVp-S219D | Supplementary Sheet | 72 | 100 |
| R1-1 | TEVp-S219D | Supplementary Sheet | 68 | 75 |
| R1-2 | TEVp-S219D | Supplementary Sheet | 69 | 73 |
| R1-3 | TEVp-S219D | Supplementary Sheet | 69 | 76 |
| R1-4 | TEVp-S219D | Supplementary Sheet | 68 | 77 |
| R1-5 | TEVp-S219D | Supplementary Sheet | 69 | 75 |
| R2-1 | R1-4 | V155L/S192N | 69 | 77 |
| R2-2 | R2-1 | T108K/I131T | 69 | 78 |
| R2-3 | R2-2 | P5F/I60K | 70 | 78 |
| R2-4 | R2-3 | I24T/L1G | 70 | 79 |
| R2-5 | R2-4 | S197H/I189L | 71 | 79 |
| R3-1 | R2-2 | R86E/F40Y | 68 | 78 |
| R3-2 | R2-2 | F20E/I24E | 70 | 78 |
| R3-3 | R2-2 | I166L/F40Y | 68 | 77 |
| R3-4 | R2-2 | M187E/S132E | 69 | 77 |
| R3-5 | R2-2 | R86E/I166L | 68 | 78 |
| R3-6 | R2-2 | I60N/I24E | 70 | 78 |
| R3-7 | R2-2 | N67E/F40Y | 69 | 78 |
| R3-8 | R2-2 | R86E/S132E | 69 | 78 |
| R3-9 | R2-2 | P73E/T22E | 69 | 77 |
| R3-10 | R2-2 | I166L/N67E | 69 | 77 |
| R3-11 | R2-2 | T191H/K58E | 69 | 77 |
| R3-12 | R2-2 | K71E/F40Y | 69 | 78 |
| R3-13 | R2-2 | S197K/E96T/I63T | 69 | 79 |
| R3-14 | R2-2 | R86E/E96T/F40L | 68 | 79 |
| R3-15 | R2-2 | S197K/R86E/I63T | 69 | 79 |
| R3-16 | R2-2 | I60K/R122T/L45A | 69 | 79 |
| R3-17 | R2-2 | S197K/I63T | 69 | 79 |
| R3-18 | R2-2 | R86E/E96T/L45A | 69 | 79 |
| R3-19 | R2-2 | S197K/N16D/E96T | 69 | 79 |
| R3-20 | R2-2 | S197K/R86E/L45A | 69 | 79 |
| R3-21 | R2-2 | F40L/F20L | 69 | 79 |
| R3-22 | R2-2 | I60K/N67S/L45A | 69 | 79 |
| R3-23 | R2-2 | S197K/R86E/L45R | 69 | 79 |
| R3-24 | R2-2 | I60K/S197K/F20Y | 69 | 79 |
| R3-25 | R2-2 | S197K/R104V/R122K | 69 | 79 |
| R3-26 | R2-2 | S197K/F40Y/L45R | 69 | 78 |
| R3-27 | R2-2 | E96T/F40Y/F20Y | 69 | 78 |
| R3-28 | R2-2 | F40Y/R86Q/F20Y | 68 | 78 |
| R3-29 | R2-2 | S197K/R86E/F40Y | 68 | 79 |
| R3-30 | R2-2 | S197K/R104V/L45R | 69 | 79 |
| R3-31 | R2-2 | S197K/E96T/L45R | 69 | 79 |
| R3-32 | R2-2 | I60K/N67E/I63E | 69 | 78 |

**Supplementary Table 2.** Production yield of TEV protease variants in *E.coli* BL21 (DE3)

| **R3 variants** | **Concentration of purified enzyme (mg/mL)** | **Yield (mg/L)** | **Fold change compared to S219D** |
| --- | --- | --- | --- |
| 1 | 1.8 | 39.2 | 8.8 |
| 2 | 3.4 | 95.8 | 21.4 |
| 3 | 1.8 | 65.5 | 14.6 |
| 4 | 1.6 | 68.4 | 15.3 |
| 5 | 1.5 | 51.1 | 11.4 |
| 6 | 2.4 | 76.5 | 17.1 |
| 7 | 1.8 | 65.6 | 14.7 |
| 8 | 3.0 | 61.7 | 13.8 |
| 9 | 2.6 | 84.4 | 18.9 |
| 10 | 0.6 | 18.7 | 4.2 |
| 11 | 2.3 | 64.4 | 14.4 |
| 12 | 2.7 | 67.2 | 15.0 |
| 13 | 2.5 | 31.5 | 7.0 |
| 14 | 4.1 | 69.2 | 15.5 |
| 15 | 5.2 | 97.5 | 21.8 |
| 16 | 5.0 | 106.4 | 23.8 |
| 17 | 2.4 | 53.3 | 11.9 |
| 18 | 3.3 | 80.9 | 18.1 |
| 19 | 2.9 | 62.0 | 13.9 |
| 20 | 4.8 | 97.6 | 21.8 |
| 21 | 4.4 | 83.1 | 18.5 |
| 22 | 2.7 | 93.1 | 20.8 |
| 23 | 6.5 | 99.0 | 22.1 |
| 24 | 2.7 | 33.7 | 7.5 |
| 25 | 6.3 | 66.2 | 14.8 |
| 26 | 3.2 | 50.8 | 11.4 |
| 27 | 2.8 | 61.2 | 13.7 |
| 28 | 4.1 | 89.7 | 20.1 |
| 29 | 5.0 | 112.7 | 25.2 |
| 30 | 3.0 | 75.6 | 16.9 |
| 31 | 2.8 | 48.8 | 10.9 |
| 32 | 5.2 | 84.2 | 18.8 |
| TEVp-S219D | 0.7 | 4.5 | 1 |
| HyperTEV60 | 6.7 | 101.0 | 22.6 |

