## Supplementary Sheet for "Combinatorial Protein Language Model–Guided Engineering of TEV Protease for Enhanced Stability and Production"

| TEV protease designs | Amino acid sequence |
|---|---|
| TEVp-S219D | GESLFKGPRDYNPISSTICHLTNESDGHTTSLYGIGFGPFIITNKHLFRRNNGTLLVQSLHGVFKVKNTTTLQQHLIDGRDMI<br>IIRMPKDFPPFPQKLKFREPQREERICLVTTNFQTKSMSSMVSDTSCTFPSSDGIFWKHWIQTKDGQCGSPLVSTRDGFIV<br>GIHSASNFTNTNNYFTSVPKNFMELLTNQEAQQWVSGWRLNADSVLWGGHKVFMDKP |
| R1-1 | LESLPPGPRDYNPIAANIVWLINTSDGYVLSLFGIGFGNDIITNRHLFWRNNGTLTITSLHGTFVVENTTTLKMHLIADRDLIII<br>RLPKDFPPFPTSLRFRVPVEGEEIVLVTRNFQTKVPRSLVSDVSRIYPSSDYTFWKHWIPTKDGQCGSPLVSVNDGTIVGLH<br>SASNFTNTNNYFMAVPENFEEVLTDEDLRRWISGWHLNSNSVMWGGHKVFMDKP |
| R1-2 | PESLPPGPRDYNPIADSIVELINTSDGFSLSLYGIGFGDYIITNAHLFWLNNGTLTVKSWHGTFLVDNTTELPVHLIPNRDLVI<br>IRLPKDFPPFPEDLRFRPPVPNESIVLVTRNFQEKPPQSVVSDVSRIRPSSNGTFWKHWIDTKDGQCGSPLVSVSDGAIVG<br>IHSASNFTNTNNYFVAVPERFMEILTNDELQNWISGWALNADSVFWGGHKVFMDKP |
| R1-3 | DESGPPGPRDYNPIADNIVLLTNISDGLRISLFGIGFGDFIITNRHLFWRNNGTLTVTSLHGTFRVPNTTDLKMHLIEDRDIVI<br>IKMPKDFPPFPTTLIFREPRANESIVLVTRNFQTKPPTSIVSDASTITPSSDGTFWKHWIETKDGQCGSPYVSVEDGTIVGLH<br>SASNFTNTNNYFVAVPVNFMRILTNEALRQWESGWHLNSYSVYWGGHKVFMDKP |
| R1-4 | LESDPPGPRDYNPIANNIVFLTNISDGYSISLFGIGFGRFIITNLHLFWRNNGTLTIKSIHGIFTINNTTKLPQHLIENRDLVIIR<br>MPKDFPPFPETLRFRTPREGETIVLVTRNFQEKTPRSVVSDVSTISPSSNGIFWKHWIPTKDGQCGSPVVSVEDGSIVGIHS<br>ASNFTNTNNYFTAVPPNFMDILTSDALRSWQSGWHLNSNSVEWGGHKVFMDKP |
| R1-5 | SESAPPGPRDYNPISNNIILLVNTSDGYRISLYGIGFGGDIITNKHLFWFNNGTLTITSIHGTFVVANTTELPMHLIPNRDLVII<br>RLPKDFPPFPEHLRFRVPREGERIVMVTRNFQTKPAISEVSDRSTITPSSNGIFWKHWIPTKDGQCGSPLVSTQDGSIVGIH<br>SASNFTNTNNYFVAVPPNFGMILTNAALRSWISGWRLNSNSVEWGGHKVFMDKP |
| R2-1 | LESDPPGPRDYNPIANNIVFLTNISDGYSISLFGIGFGRFIITNLHLFWRNNGTLTIKSIHGIFTINNTTKLPQHLIENRDLVIIR<br>MPKDFPPFPETLRFRTPREGETIVLVTRNFQEKTPRSVVSDVSTISPSSNGIFWKHWIPTKDGQCGSPLVSVEDGSIVGIHS<br>ASNFTNTNNYFTAVPPNFMDILTNDALRSWQSGWHLNSNSVEWGGHKVFMDKP |
| R2-2 | LESDPPGPRDYNPIANNIVFLTNISDGYSISLFGIGFGRFIITNLHLFWRNNGTLTIKSIHGIFTINNTTKLPQHLIENRDLVIIR<br>MPKDFPPFPETLRFRTPREGEKIVLVTRNFQEKTPRSVVSDVSTTSPSSNGIFWKHWIPTKDGQCGSPLVSVEDGSIVGIHS<br>ASNFTNTNNYFTAVPPNFMDILTNDALRSWQSGWHLNSNSVEWGGHKVFMDKP |
| R2-3 | LESDFPGPRDYNPIANNIVFLTNISDGYSISLFGIGFGRFIITNLHLFWRNNGTLTIKSKHGIFTINNTTKLPQHLIENRDLVIIR<br>MPKDFPPFPETLRFRTPREGEKIVLVTRNFQEKTPRSVVSDVSTTSPSSNGIFWKHWIPTKDGQCGSPLVSVEDGSIVGIHS<br>ASNFTNTNNYFTAVPPNFMDILTNDALRSWQSGWHLNSNSVEWGGHKVFMDKP |
| R2-4 | GESDFPGPRDYNPIANNIVFLTNTSDGYSISLFGIGFGRFIITNLHLFWRNNGTLTIKSKHGIFTINNTTKLPQHLIENRDLVII<br>RMPKDFPPFPETLRFRTPREGEKIVLVTRNFQEKTPRSVVSDVSTTSPSSNGIFWKHWIPTKDGQCGSPLVSVEDGSIVGIH<br>SASNFTNTNNYFTAVPPNFMDILTNDALRSWQSGWHLNSNSVEWGGHKVFMDKP |
| R2-5 | GESDFPGPRDYNPIANNIVFLTNTSDGYSISLFGIGFGRFIITNLHLFWRNNGTLTIKSKHGIFTINNTTKLPQHLIENRDLVII<br>RMPKDFPPFPETLRFRTPREGEKIVLVTRNFQEKTPRSVVSDVSTTSPSSNGIFWKHWIPTKDGQCGSPLVSVEDGSIVGIH<br>SASNFTNTNNYFTAVPPNFMDLLTNDALRHWQSGWHLNSNSVEWGGHKVFMDKP |
| R3-1 | LESDPPGPRDYNPIANNIVFLTNISDGYSISLFGIGFGRYIITNLHLFWRNNGTLTIKSIHGIFTINNTTKLPQHLIENRDLVIIE<br>MPKDFPPFPETLRFRTPREGEKIVLVTRNFQEKTPRSVVSDVSTTSPSSNGIFWKHWIPTKDGQCGSPLVSVEDGSIVGIHS<br>ASNFTNTNNYFTAVPPNFMDILTNDALRSWQSGWHLNSNSVEWGGHKVFMDKP |
| R3-2 | LESDPPGPRDYNPIANNIVELTNESDGYSISLFGIGFGRFIITNLHLFWRNNGTLTIKSIHGIFTINNTTKLPQHLIENRDLVIIR<br>MPKDFPPFPETLRFRTPREGEKIVLVTRNFQEKTPRSVVSDVSTTSPSSNGIFWKHWIPTKDGQCGSPLVSVEDGSIVGIHS<br>ASNFTNTNNYFTAVPPNFMDILTNDALRSWQSGWHLNSNSVEWGGHKVFMDKP |
| R3-3 | LESDPPGPRDYNPIANNIVFLTNISDGYSISLFGIGFGRYIITNLHLFWRNNGTLTIKSIHGIFTINNTTKLPQHLIENRDLVIIR<br>MPKDFPPFPETLRFRTPREGEKIVLVTRNFQEKTPRSVVSDVSTTSPSSNGIFWKHWIPTKDGQCGSPLVSVEDGSIVGLH<br>SASNFTNTNNYFTAVPPNFMDILTNDALRSWQSGWHLNSNSVEWGGHKVFMDKP |
| R3-4 | LESDPPGPRDYNPIANNIVFLTNISDGYSISLFGIGFGRFIITNLHLFWRNNGTLTIKSIHGIFTINNTTKLPQHLIENRDLVIIR<br>MPKDFPPFPETLRFRTPREGEKIVLVTRNFQEKTPRSVVSDVSTTEPSSNGIFWKHWIPTKDGQCGSPLVSVEDGSIVGIHS<br>ASNFTNTNNYFTAVPPNFEDILTNDALRSWQSGWHLNSNSVEWGGHKVFMDKP |
| R3-5 | LESDPPGPRDYNPIANNIVFLTNISDGYSISLFGIGFGRFIITNLHLFWRNNGTLTIKSIHGIFTINNTTKLPQHLIENRDLVIIE<br>MPKDFPPFPETLRFRTPREGEKIVLVTRNFQEKTPRSVVSDVSTTSPSSNGIFWKHWIPTKDGQCGSPLVSVEDGSIVGLH<br>SASNFTNTNNYFTAVPPNFMDILTNDALRSWQSGWHLNSNSVEWGGHKVFMDKP |
| R3-6 | LESDPPGPRDYNPIANNIVFLTNESDGYSISLFGIGFGRFIITNLHLFWRNNGTLTIKSNHGIFTINNTTKLPQHLIENRDLVII<br>RMPKDFPPFPETLRFRTPREGEKIVLVTRNFQEKTPRSVVSDVSTTSPSSNGIFWKHWIPTKDGQCGSPLVSVEDGSIVGIH<br>SASNFTNTNNYFTAVPPNFMDILTNDALRSWQSGWHLNSNSVEWGGHKVFMDKP |
| R3-7 | LESDPPGPRDYNPIANNIVFLTNISDGYSISLFGIGFGRYIITNLHLFWRNNGTLTIKSIHGIFTIENTTKLPQHLIENRDLVIIRM<br>PKDFPPFPETLRFRTPREGEKIVLVTRNFQEKTPRSVVSDVSTTSPSSNGIFWKHWIPTKDGQCGSPLVSVEDGSIVGIHSA<br>SNFTNTNNYFTAVPPNFMDILTNDALRSWQSGWHLNSNSVEWGGHKVFMDKP |
| R3-8 | LESDPPGPRDYNPIANNIVFLTNISDGYSISLFGIGFGRFIITNLHLFWRNNGTLTIKSIHGIFTINNTTKLPQHLIENRDLVIIE<br>MPKDFPPFPETLRFRTPREGEKIVLVTRNFQEKTPRSVVSDVSTTEPSSNGIFWKHWIPTKDGQCGSPLVSVEDGSIVGIHS<br>ASNFTNTNNYFTAVPPNFMDILTNDALRSWQSGWHLNSNSVEWGGHKVFMDKP |

| | |
|---|---|
| R3-9 | LESDPPGPRDYNPIANNIVFLENISDGYSISLFGIGFGRFIITNLHLFWRNNGTLTIKSIHGIFTINNTTKLEQHLIENRDLVIIR<br>MPKDFPPFPETLRFRTPREGEKIVLVTRNFQEKTPRSVVSDVSTTSPSSNGIFWKHWIPTKDGQCGSPLVSVEDGSIVGIHS<br>ASNFTNTNNYFTAVPPNFMDILTNDALRSWQSGWHLNSNSVEWGGHKVFMDKP |
| R3-10 | LESDPPGPRDYNPIANNIVFLTNISDGYSISLFGIGFGRFIITNLHLFWRNNGTLTIKSIHGIFTIENTTKLPQHLIENRDLVIIRM<br>PKDFPPFPETLRFRTPREGEKIVLVTRNFQEKTPRSVVSDVSTTSPSSNGIFWKHWIPTKDGQCGSPLVSVEDGSIVGLHSA<br>SNFTNTNNYFTAVPPNFMDILTNDALRSWQSGWHLNSNSVEWGGHKVFMDKP |
| R3-11 | LESDPPGPRDYNPIANNIVFLTNISDGYSISLFGIGFGRFIITNLHLFWRNNGTLTIESIHGIFTINNTTKLPQHLIENRDLVIIR<br>MPKDFPPFPETLRFRTPREGEKIVLVTRNFQEKTPRSVVSDVSTTSPSSNGIFWKHWIPTKDGQCGSPLVSVEDGSIVGIHS<br>ASNFTNTNNYFTAVPPNFMDILHNDALRSWQSGWHLNSNSVEWGGHKVFMDKP |
| R3-12 | LESDPPGPRDYNPIANNIVFLTNISDGYSISLFGIGFGRYIITNLHLFWRNNGTLTIKSIHGIFTINNTTELPQHLIENRDLVIIR<br>MPKDFPPFPETLRFRTPREGEKIVLVTRNFQEKTPRSVVSDVSTTSPSSNGIFWKHWIPTKDGQCGSPLVSVEDGSIVGIHS<br>ASNFTNTNNYFTAVPPNFMDILTNDALRSWQSGWHLNSNSVEWGGHKVFMDKP |
| R3-13 | LESDPPGPRDYNPIANNIVFLTNISDGYSISLFGIGFGRFIITNLHLFWRNNGTLTIKSIHGTFTINNTTKLPQHLIENRDLVIIR<br>MPKDFPPFPTTLRFRTPREGEKIVLVTRNFQEKTPRSVVSDVSTTSPSSNGIFWKHWIPTKDGQCGSPLVSVEDGSIVGIHS<br>ASNFTNTNNYFTAVPPNFMDILTNDALRKWQSGWHLNSNSVEWGGHKVFMDKP |
| R3-14 | LESDPPGPRDYNPIANNIVFLTNISDGYSISLFGIGFGRLIITNLHLFWRNNGTLTIKSIHGIFTINNTTKLPQHLIENRDLVIIE<br>MPKDFPPFPTTLRFRTPREGEKIVLVTRNFQEKTPRSVVSDVSTTSPSSNGIFWKHWIPTKDGQCGSPLVSVEDGSIVGIHS<br>ASNFTNTNNYFTAVPPNFMDILTNDALRSWQSGWHLNSNSVEWGGHKVFMDKP |
| R3-15 | LESDPPGPRDYNPIANNIVFLTNISDGYSISLFGIGFGRFIITNLHLFWRNNGTLTIKSIHGTFTINNTTKLPQHLIENRDLVIIE<br>MPKDFPPFPETLRFRTPREGEKIVLVTRNFQEKTPRSVVSDVSTTSPSSNGIFWKHWIPTKDGQCGSPLVSVEDGSIVGIHS<br>ASNFTNTNNYFTAVPPNFMDILTNDALRKWQSGWHLNSNSVEWGGHKVFMDKP |
| R3-16 | LESDPPGPRDYNPIANNIVFLTNISDGYSISLFGIGFGRFIITNAHLFWRNNGTLTIKSKHGIFTINNTTKLPQHLIENRDLVIIR<br>MPKDFPPFPETLRFRTPREGEKIVLVTRNFQEKTPTSVVSDVSTTSPSSNGIFWKHWIPTKDGQCGSPLVSVEDGSIVGIHS<br>ASNFTNTNNYFTAVPPNFMDILTNDALRSWQSGWHLNSNSVEWGGHKVFMDKP |
| R3-17 | LESDPPGPRDYNPIANNIVFLTNISDGYSISLFGIGFGRFIITNLHLFWRNNGTLTIKSIHGTFTINNTTKLPQHLIENRDLVIIR<br>MPKDFPPFPETLRFRTPREGEKIVLVTRNFQEKTPRSVVSDVSTTSPSSNGIFWKHWIPTKDGQCGSPLVSVEDGSIVGIHS<br>ASNFTNTNNYFTAVPPNFMDILTNDALRKWQSGWHLNSNSVEWGGHKVFMDKP |
| R3-18 | LESDPPGPRDYNPIANNIVFLTNISDGYSISLFGIGFGRFIITNAHLFWRNNGTLTIKSIHGIFTINNTTKLPQHLIENRDLVIIE<br>MPKDFPPFPTTLRFRTPREGEKIVLVTRNFQEKTPRSVVSDVSTTSPSSNGIFWKHWIPTKDGQCGSPLVSVEDGSIVGIHS<br>ASNFTNTNNYFTAVPPNFMDILTNDALRSWQSGWHLNSNSVEWGGHKVFMDKP |
| R3-19 | LESDPPGPRDYNPIADNIVFLTNISDGYSISLFGIGFGRFIITNLHLFWRNNGTLTIKSIHGIFTINNTTKLPQHLIENRDLVIIR<br>MPKDFPPFPTTLRFRTPREGEKIVLVTRNFQEKTPRSVVSDVSTTSPSSNGIFWKHWIPTKDGQCGSPLVSVEDGSIVGIHS<br>ASNFTNTNNYFTAVPPNFMDILTNDALRKWQSGWHLNSNSVEWGGHKVFMDKP |
| R3-20 | LESDPPGPRDYNPIANNIVFLTNISDGYSISLFGIGFGRFIITNAHLFWRNNGTLTIKSIHGIFTINNTTKLPQHLIENRDLVIIE<br>MPKDFPPFPETLRFRTPREGEKIVLVTRNFQEKTPRSVVSDVSTTSPSSNGIFWKHWIPTKDGQCGSPLVSVEDGSIVGIHS<br>ASNFTNTNNYFTAVPPNFMDILTNDALRKWQSGWHLNSNSVEWGGHKVFMDKP |
| R3-21 | LESDPPGPRDYNPIANNIVLLTNISDGYSISLFGIGFGRLIITNLHLFWRNNGTLTIKSIHGIFTINNTTKLPQHLIENRDLVIIR<br>MPKDFPPFPETLRFRTPREGEKIVLVTRNFQEKTPRSVVSDVSTTSPSSNGIFWKHWIPTKDGQCGSPLVSVEDGSIVGIHS<br>ASNFTNTNNYFTAVPPNFMDILTNDALRSWQSGWHLNSNSVEWGGHKVFMDKP |
| R3-22 | LESDPPGPRDYNPIANNIVFLTNISDGYSISLFGIGFGRFIITNAHLFWRNNGTLTIKSKHGIFTISNTTKLPQHLIENRDLVIIR<br>MPKDFPPFPETLRFRTPREGEKIVLVTRNFQEKTPRSVVSDVSTTSPSSNGIFWKHWIPTKDGQCGSPLVSVEDGSIVGIHS<br>ASNFTNTNNYFTAVPPNFMDILTNDALRSWQSGWHLNSNSVEWGGHKVFMDKP |
| R3-23 | LESDPPGPRDYNPIANNIVFLTNISDGYSISLFGIGFGRFIITNRHLFWRNNGTLTIKSIHGIFTINNTTKLPQHLIENRDLVIIE<br>MPKDFPPFPETLRFRTPREGEKIVLVTRNFQEKTPRSVVSDVSTTSPSSNGIFWKHWIPTKDGQCGSPLVSVEDGSIVGIHS<br>ASNFTNTNNYFTAVPPNFMDILTNDALRKWQSGWHLNSNSVEWGGHKVFMDKP |
| R3-24 | LESDPPGPRDYNPIANNIVYLTNISDGYSISLFGIGFGRFIITNLHLFWRNNGTLTIKSKHGIFTINNTTKLPQHLIENRDLVIIR<br>MPKDFPPFPETLRFRTPREGEKIVLVTRNFQEKTPRSVVSDVSTTSPSSNGIFWKHWIPTKDGQCGSPLVSVEDGSIVGIHS<br>ASNFTNTNNYFTAVPPNFMDILTNDALRKWQSGWHLNSNSVEWGGHKVFMDKP |
| R3-25 | LESDPPGPRDYNPIANNIVFLTNISDGYSISLFGIGFGRFIITNLHLFWRNNGTLTIKSIHGIFTINNTTKLPQHLIENRDLVIIR<br>MPKDFPPFPETLRFRTPVEGEKIVLVTRNFQEKTPKSVVSDVSTTSPSSNGIFWKHWIPTKDGQCGSPLVSVEDGSIVGIHS<br>ASNFTNTNNYFTAVPPNFMDILTNDALRKWQSGWHLNSNSVEWGGHKVFMDKP |
| R3-26 | LESDPPGPRDYNPIANNIVFLTNISDGYSISLFGIGFGRYIITNRHLFWRNNGTLTIKSIHGIFTINNTTKLPQHLIENRDLVIIR<br>MPKDFPPFPETLRFRTPREGEKIVLVTRNFQEKTPRSVVSDVSTTSPSSNGIFWKHWIPTKDGQCGSPLVSVEDGSIVGIHS<br>ASNFTNTNNYFTAVPPNFMDILTNDALRKWQSGWHLNSNSVEWGGHKVFMDKP |
| R3-27 | LESDPPGPRDYNPIANNIVYLTNISDGYSISLFGIGFGRYIITNLHLFWRNNGTLTIKSIHGIFTINNTTKLPQHLIENRDLVIIR<br>MPKDFPPFPTTLRFRTPREGEKIVLVTRNFQEKTPRSVVSDVSTTSPSSNGIFWKHWIPTKDGQCGSPLVSVEDGSIVGIHS<br>ASNFTNTNNYFTAVPPNFMDILTNDALRSWQSGWHLNSNSVEWGGHKVFMDKP |
| R3-28 | LESDPPGPRDYNPIANNIVYLTNISDGYSISLFGIGFGRYIITNLHLFWRNNGTLTIKSIHGIFTINNTTKLPQHLIENRDLVIIQ<br>MPKDFPPFPETLRFRTPREGEKIVLVTRNFQEKTPRSVVSDVSTTSPSSNGIFWKHWIPTKDGQCGSPLVSVEDGSIVGIHS<br>ASNFTNTNNYFTAVPPNFMDILTNDALRSWQSGWHLNSNSVEWGGHKVFMDKP |

| | |
|---|---|
| R3-29 | LESDPPGPRDYNPIANNIVFLTNISDGYSISLFGIGFGRYIITNLHLFWRNNGTLTIKSIHGIFTINNTTKLPQHLIENRDLVIIE<br>MPKDFPPFPETLRFRTPREGEKIVLVTRNFQEKTPRSVVSDVSTTSPSSNGIFWKHWIPTKDGQCGSPLVSVEDGSIVGIHS<br>ASNFTNTNNYFTAVPPNFMDILTNDALRKWQSGWHLNSNSVEWGGHKVFMDKP |
| R3-30 | LESDPPGPRDYNPIANNIVFLTNISDGYSISLFGIGFGRFIITNRHLFWRNNGTLTIKSIHGIFTINNTTKLPQHLIENRDLVIIR<br>MPKDFPPFPETLRFRTPVEGEKIVLVTRNFQEKTPRSVVSDVSTTSPSSNGIFWKHWIPTKDGQCGSPLVSVEDGSIVGIHS<br>ASNFTNTNNYFTAVPPNFMDILTNDALRKWQSGWHLNSNSVEWGGHKVFMDKP |
| R3-31 | LESDPPGPRDYNPIANNIVFLTNISDGYSISLFGIGFGRFIITNRHLFWRNNGTLTIKSIHGIFTINNTTKLPQHLIENRDLVIIR<br>MPKDFPPFPTTLRFRTPREGEKIVLVTRNFQEKTPRSVVSDVSTTSPSSNGIFWKHWIPTKDGQCGSPLVSVEDGSIVGIHS<br>ASNFTNTNNYFTAVPPNFMDILTNDALRKWQSGWHLNSNSVEWGGHKVFMDKP |
| R3-32 | LESDPPGPRDYNPIANNIVFLTNISDGYSISLFGIGFGRFIITNLHLFWRNNGTLTIKSKHGEFTIENTTKLPQHLIENRDLVIIR<br>MPKDFPPFPETLRFRTPREGEKIVLVTRNFQEKTPRSVVSDVSTTSPSSNGIFWKHWIPTKDGQCGSPLVSVEDGSIVGIHS<br>ASNFTNTNNYFTAVPPNFMDILTNDALRSWQSGWHLNSNSVEWGGHKVFMDKP |
| hyperTEV60 | AESAAPGPRDYNPISDTIVLLTNTSDGYSISLYGIGFGPLIITNAHLFRRNNGTLTITSKHGTFTISNTTTLKLHLIEGRDLVLIEM<br>PKDFPPFPTNLVFREPVVGEEIVLVTRNFQTKTPTSEVSDVSTTYPSSDGVFWKHWIPTKDGQCGSPMVSVTDGSIVGIHS<br>ASNFTNTNNYFTAVPPDFMRLLTDPSLQKWVSGWSLNSDSVEWGGHKVFMDKP |
